# Phenotyping stomatal closure by thermal imaging for GWAS and TWAS of water use efficiency-related genes

**DOI:** 10.1101/2021.05.06.442962

**Authors:** Charles P. Pignon, Samuel B. Fernandes, Ravi Valluru, Nonoy Bandillo, Roberto Lozano, Edward Buckler, Michael A. Gore, Stephen P. Long, Patrick J. Brown, Andrew D. B. Leakey

**Affiliations:** Department of Plant Biology, University of Illinois at Urbana-Champaign, Urbana, IL 61801, USA; Department of Crop Sciences, University of Illinois at Urbana-Champaign, Urbana, IL 61801, USA; Carl R. Woese Institute for Genomic Biology, University of Illinois at Urbana-Champaign, Urbana, IL 61801, USA; Institute for Genomic Diversity, Cornell University, Ithaca, NY 14853, USA; Lincoln Institute for Agri-Food Technology, University of Lincoln, LN1 3QE Lincoln, UK; Department of Plant Sciences, North Dakota State University, Fargo, ND 58105, USA; Plant Breeding and Genetics Section, School of Integrative Plant Science, Cornell University, Ithaca, NY 14853, USA; United States Department of Agriculture, Agricultural Research Service (USDA-ARS) R.W. Holley Center for Agriculture and Health, Ithaca, NY 14853, USA; Lancaster Environment Centre, University of Lancaster, LA1 1YX, UK

## Abstract

Stomata allow CO_2_ uptake by leaves for photosynthetic assimilation at the cost of water vapor loss to the atmosphere. The opening and closing of stomata in response to fluctuations in light intensity regulate CO_2_ and water fluxes and are essential to maintenance of water-use efficiency (WUE). However, little is known about the genetic basis for natural variation in stomatal movement, especially in C_4_ crops. This is partly because the stomatal response to a change in light intensity is difficult to measure at the scale required for association studies. High-throughput thermal imaging was used to bypass the phenotyping bottleneck and assess 10 traits describing stomatal conductance (*g_s_*) before, during and after a stepwise decrease in light intensity for a diversity panel of 659 sorghum accessions. Results from thermal imaging significantly correlated with photosynthetic gas-exchange measurements. *g_s_* traits varied substantially across the population and were moderately heritable (*h^2^* up to 0.72). An integrated genome-wide and transcriptome-wide association study (GWAS/TWAS) identified candidate genes putatively driving variation in stomatal conductance traits. Of the 239 unique candidate genes identified with greatest confidence, 77 were orthologs of Arabidopsis genes related to functions implicated in WUE, including stomatal opening/closing (24 genes), stomatal/epidermal cell development (35 genes), leaf/vasculature development (12 genes), or chlorophyll metabolism/photosynthesis (8 genes). These findings demonstrate an approach to finding genotype-to-phenotype relationships for a challenging trait as well as candidate genes for further investigation of the genetic basis of WUE in a model C_4_ grass for bioenergy, food, and forage production.

**One sentence summary:** Rapid phenotyping of 659 accessions of *Sorghum bicolor* revealed heritable stomatal responses to a decrease in light. GWAS/TWAS was used to identify candidate genes influencing traits important to WUE.

## Introduction

Water availability is a major limiting factor to agriculture worldwide (Boyer, 1982), and is predicted to become even more limiting due to rising demand for water resulting from increasing atmospheric vapor pressure deficit (VPD) with climate change (Lobell et al., 2008; WWAP, 2015; FAO et al., 2018). Greater crop water use associated with greater above-ground biomass has also been implicated in past and future increases in crop yield (Sinclair et al., 1984; Ray et al., 2013; Lobell et al., 2014; Ort and Long, 2014; Koester et al. 2016; DeLucia et al., 2019). Therefore, unless WUE is enhanced, agricultural systems in the near future will be increasingly threatened by drought, and unsustainable practices such as over-irrigation may result (DeLucia et al., 2019; Leakey et al., 2019).

WUE is key to terrestrial plant growth due to the inherent trade-off between net photosynthetic carbon assimilation (*A*) and water loss through transpiration (Wong et al., 1979). Most leaf water and CO_2_ fluxes pass through stomatal pores (Kerstiens, 1996; Hetherington and Woodward, 2003). Stomatal aperture is dynamically regulated by the movement of stomatal guard cells (Assmann and Jegla, 2016), which respond to extrinsic and intrinsic signals to optimize CO_2_ uptake relative to water vapor loss (Medlyn et al 2011). The synchrony of stomatal opening and closing with fluctuations in photosynthetic CO_2_ assimilation varies within and among species, with significant consequences for intrinsic water-use efficiency (*iWUE*, the ratio of *A* to stomatal conductance to water vapor (*g_s_*)) (Lawson and Blatt, 2014; Kaiser et al., 2015). If conditions are favorable for *A,* but *g_s_* is low, *A* will be limited by CO_2_ supply. Conversely, if *A* is low, but *g_s_* is high, unnecessary transpiration will occur with no corresponding benefit to *A*, substantially decreasing *iWUE*. This is important in fluctuating light environments such as field crop canopies, where the movement of clouds and leaves cause frequent and abrupt changes in photosynthetic photon flux density (PPFD) at the leaf level (Pearcy, 1990; Zhu et al., 2004; Way and Pearcy, 2012; Wang et al., 2020).

Generally, *g_s_* responds an order of magnitude more slowly than *A* to a decline in PPFD, declining to a new steady-state over the course of several minutes (Hetherington and Woodward, 2003; McAusland et al., 2016). This results in a de-synchronization of *A* and *g_s_* and loss of *iWUE*. Ensuring that *A* and *g_s_* are synchronized in their response to fluctuating light, by accelerating stomatal movement, could yield substantial benefits to *iWUE* of 20-30% (Lawson and Blatt, 2014). Proof-of-concept from transgenic manipulation of the stomatal light-sensing mechanism has led to accelerated stomatal movement and improved *iWUE* in fluctuating light in Arabidopsis and tobacco (Glowacka et al., 2018; Papanatsiou et al., 2019). Still, accelerating stomatal movement by breeding or biotechnology for improved *iWUE* remains an open challenge in cereal crops (Faralli et al., 2019).

The genetic basis for natural variation in traits describing stomatal opening/closing is poorly understood, in part because such traits are difficult to phenotype at the scale required for association studies. The response of *g_s_* to fluctuating light is lengthy, with some species needing >30 minutes to transition from steady-state at one light intensity to another (Lawson and Blatt, 2014; McAusland et al., 2016; Deans et al., 2019). The standard measurement of *g_s_* using gas-exchange chambers or porometers requires a single instrument for each leaf, so is costly in terms of personnel and equipment. Using thermal imaging to track changes in leaf temperature resulting from stomatal movement, with the leaf warming up as stomata close due to a proportionate reduction of cooling by transpiration, provides a high throughput phenotyping method in which numerous leaves can be measured simultaneously (Jones et al., 2002; Guilioni et al., 2008; Vialet-Chabrand and Lawson, 2019). Thermal imaging has been used to monitor stomatal closure (Jones et al., 2002) and stress response (Grant et al., 2007) in a grapevine field, to identify Arabidopsis mutants with altered *g_s_* (Merlot et al., 2002), and coupled with chlorophyll fluorescence measurements to screen Arabidopsis plants for *iWUE* (McAusland et al., 2013). In a broad validation experiment, thermal measurements predicted *g_s_* for a range of species exposed to different experimental treatments (Grant et al., 2006). Finally, thermal imaging was used to perform high-throughput phenotyping of the response of hundreds of ecotypes of Arabidopsis to changing light and [CO_2_] (Takahashi et al., 2015). However, the ability of thermal imaging to produce trait estimates with sufficiently high heritability to support quantitative genetic investigation of genotype-to-phenotype relationships for stomatal opening/closing remains unclear.

Genome-wide association studies (GWAS) are widely used to identify the genetic basis for natural variation in agronomic, developmental, physiological and biochemical traits in plant species, including sorghum (Casa et al., 2008; Morris et al., 2013; Burks et al., 2015; Ortiz et al., 2017). The benefits and limitations of the approach have been reviewed in many contexts (Liu and Yan, 2019; Tam et al., 2019; Zhou and Huang, 2019). GWAS of stomatal movement can be challenging because biophysical trade-offs can limit the extent of natural variation in the trait, and because measurements may be slow, sensitive to environmental conditions, and lack accuracy, precision or both. Together, these factors can reduce the variance that can be ascribed to genotype, i.e. reduce heritability, and constrain the size of the mapping population that can be studied, resulting in low statistical power. Recently, combining GWAS with transcriptome-wide association study (TWAS), which identifies significant associations between trait variation and RNA transcript abundance across all expressed genes in a tissue, has been shown to improve identification of genes underlying trait variation (Kremling et al., 2019).

This study aimed to demonstrate how phenotyping by thermal imaging could be used in conjunction with integrated GWAS/TWAS to quantify natural diversity in stomatal/opening closing and identify genotype-to-phenotype relationships in a model C_4_ crop. *Sorghum bicolor* ((L.) Moench) is a model species used to study photosynthesis, abiotic/biotic stress and canopy architecture, as well as applied investigations in the context of food, fuel and forage production (Paterson et al., 2009; Morris et al., 2013). It has particular importance in semi-arid conditions due to its high productivity and drought-tolerance (Regassa and Wortmann, 2014; Hadebe et al., 2017). Sorghum is an especially interesting model in which to study stomatal movement because it has fast stomatal responses to decreasing light relative to other species (McAusland et al., 2016; Pignon et al., 2021). High-throughput thermal imaging was used to measure 10 traits describing the response of *g_s_* to a decrease in photosynthetic photon flux density (*PPFD*) in over 2000 plants of 659 accessions in a sorghum association mapping population. Results were validated against photosynthetic gas-exchange measurements. Phenotypic trait correlations were used to identify general patterns in stomatal behavior. GWAS and TWAS were performed to identify phenotype to genotype associations, along with an ensemble approach combining GWAS and TWAS results using the Fisher’s combined test (FCT) (Kremling et al., 2019), followed by a GO enrichment analysis. The resulting list of 239 candidate genes identified with greatest confidence was enriched in orthologs of genes implicated in stomatal and photosynthetic traits in Arabidopsis and maize.

## Results

The response of *g_s_* estimated from thermal imaging (*g_s thermal_*) to a reduction in *PPFD* from 750 to 75 µmol m^-2^ s^-1^ was measured in 659 sorghum accessions (Table 1; Fig. 1). On a subset of 64 plants, *g_s_* estimates were also obtained from gas-exchange measurements to validate *g_s thermal_* as a proxy for *g_s_*. Both *g_s thermal_* and *g_s_* predicted a similar pattern of stomatal closure upon a decrease in *PPFD*, sometimes followed by re-opening at low *PPFD* (Fig. 2). All traits derived from *g_s thermal_* were significantly and positively correlated with their equivalents from *g_s_* (*p*<0.005, Pearson’s *r* ranging from 0.38 - 0.59, Spearman’s rank-order ρ ranging from 0.29 - 0.66, Fig. 3).

**Figure 1:**
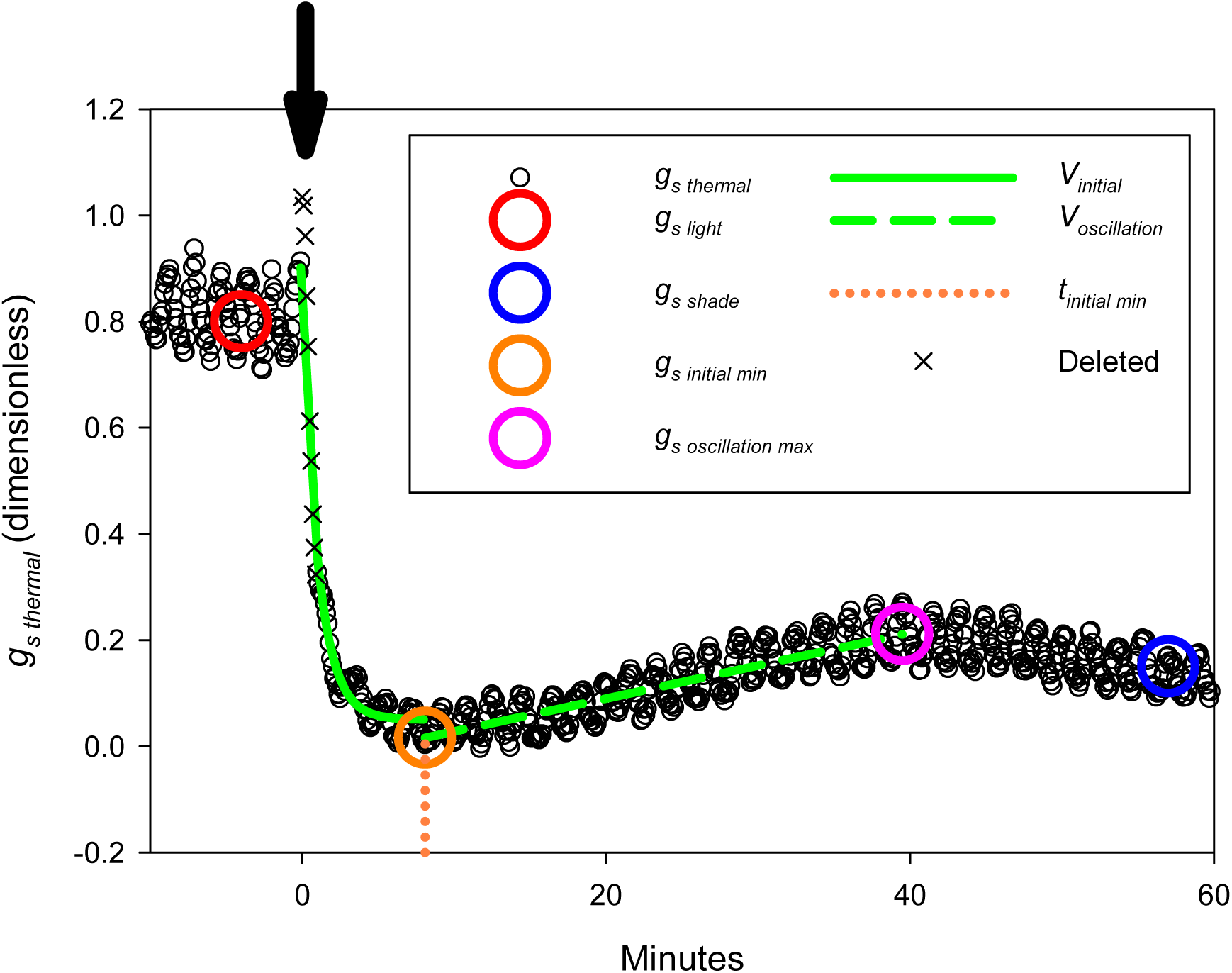
Schematic of *g_s thermal_* analysis method, where *g_s thermal_* is a proxy for stomatal conductance to water vapor (*g_s_*) that is derived from thermal imaging. Each response was measured on a single leaf of sorghum, exposed to *PPFD*=750 µmol m^-2^ s^-1^ for 40 minutes. At t=0, indicated by an arrow, *PPFD* was reduced by 90%. Black circles show *g_s thermal_*. Crosses show measurements from t=0 to t=0.9 minutes which were removed because they consistently showed an anomalous spike. *g_s light_*, the steady-state high-*PPFD* value of *g_s thermal_*, was the mean *g_s thermal_* from t=-5 to 0 minutes (red circle). *g_s shade_*, the steady-state low-*PPFD* value of *g_s thermal_*, was the mean *g_s thermal_* from t=52 to 60 minutes (blue circle). *g_s Σ shade_* was the area under the curve from t=0 to 60 minutes. *g_s initial min_* was the minimum of *g_s thermal_* reached immediately after the decrease in *PPFD* (orange circle). *g_s oscillation max_* was the maximum of *g_s thermal_* reached during stomatal re-opening at low *PPFD* (pink circle). The time at which *g_s thermal_* reached 110% of *g_s initial min_* was recorded as *t _initial min_* (dotted orange line). *V_initial_*, the initial rate of decline in *g_s thermal_* after the decrease in *PPFD*, was the exponential rate of decline of *g_s thermal_* t=- 0.1 minutes to t= *t _initial min_* (solid green line). *V_oscillation_* was the linear rate of increase in *g_s thermal_* during stomatal reopening at low *PPFD* (dashed green line).

**Figure 2:**
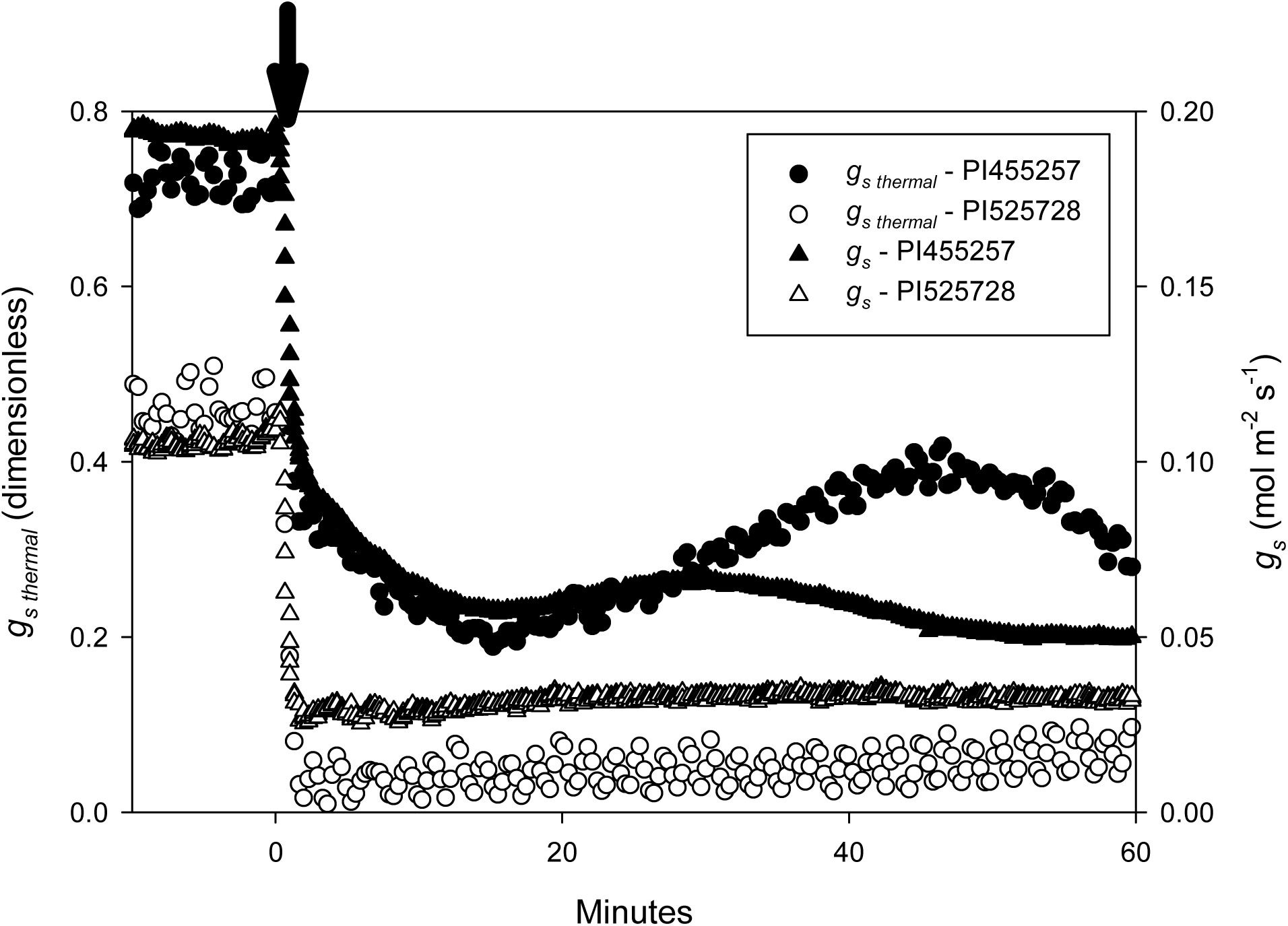
Representative responses of *g_s thermal_* and *g_s_* to a 90% drop in *PPFD* for two accessions: PI455257 and PI525728. *g_s thermal_* was measured from thermal imaging as a proxy for *g_s_*. Each *g_s thermal_* response curve was measured on a single leaf, acclimated to *PPFD*=750 µmol m^-2^ s^-1^ for 40 minutes, then *PPFD* was reduced by 90% for 60 minutes at t=0, indicated by an arrow. On a subset of plants, including the two shown here, the protocol was immediately repeated to measure *g_s_* using gas exchange.

**Figure 3:**
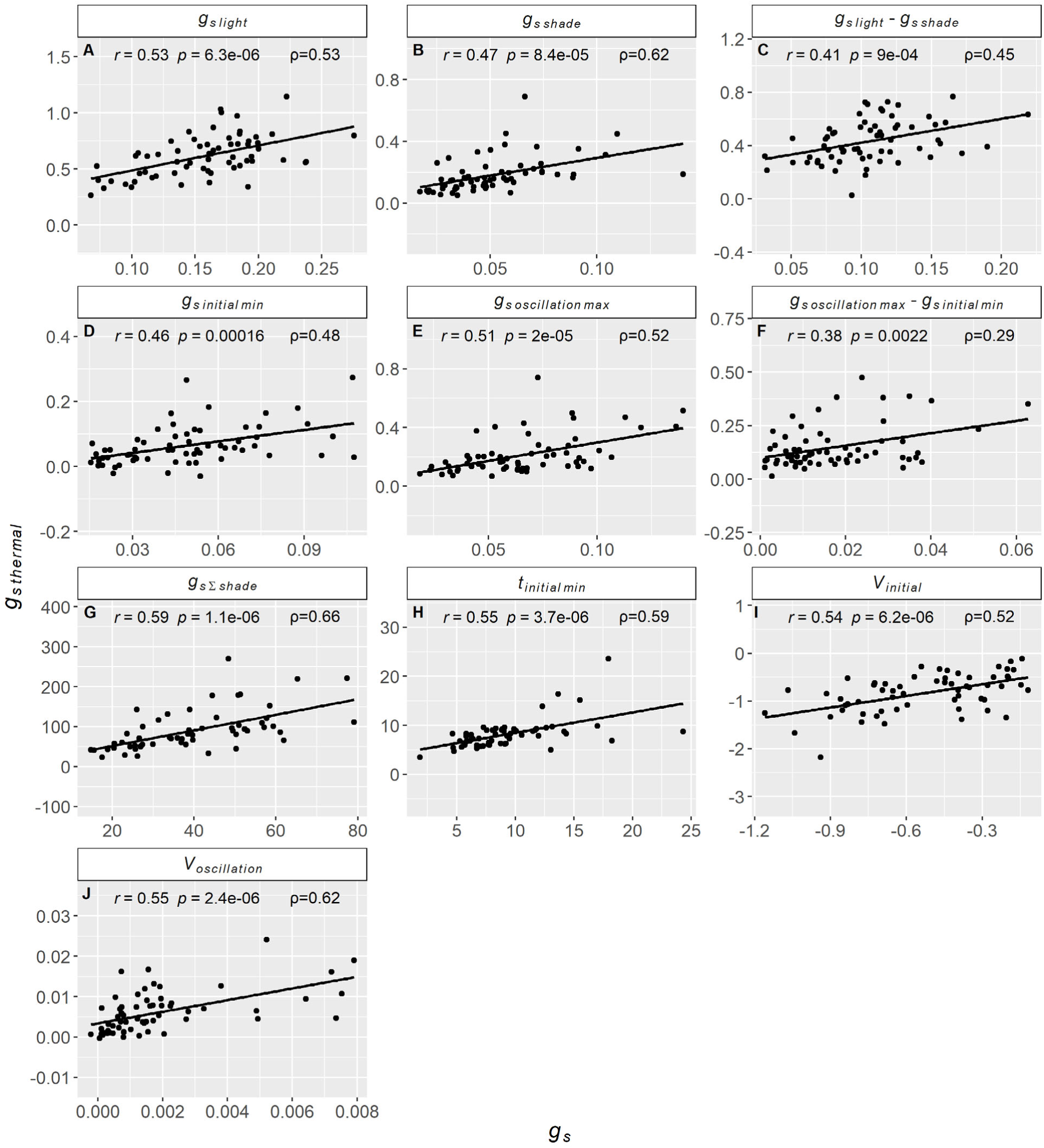
Correlation scatterplots between stomatal conductance traits derived from *g_s thermal_* measurements and their counterparts derived from *g_s_* measurements. Data are for a subset of plants that were first imaged using a thermal camera to obtain *g_s thermal_*, then immediately measured with gas exchange to obtain *g_s_*. *t_initial min_* is in minutes, all other *g_s thermal_* traits are dimensionless. Pearson’s *r* and the associated *p*-value, along with Spearman’s rank-order ρ, are also given.

**Table 1:** Descriptive statistics of stomatal conductance traits. *h^2^* is genomic heritability. *t_initial min_* is in minutes, all other traits are dimensionless. Graphical description of these traits is given in Fig. 1.

| Trait | Description | Mean | SD | Min | Max | $h^2$ |
| --- | --- | --- | --- | --- | --- | --- |
| $g_s\ light$ | Steady-state $g_s\ thermal$ at high *PPFD* | 0.58 | 0.08 | 0.31 | 0.89 | 0.31 |
| $g_s\ shade$ | Steady-state $g_s\ thermal$ at low *PPFD* | 0.17 | 0.07 | 0.05 | 0.49 | 0.70 |
| $(g_s\ light - g_s\ shade)$ | Amplitude of overall decline in $g_s\ thermal$ from high to low *PPFD* | 0.41 | 0.08 | 0.11 | 0.66 | 0.31 |
| $g_s\ initial\ min$ | Minimum $g_s\ thermal$ reached after the *PPFD* decrease | 0.05 | 0.04 | -0.03 | 0.24 | 0.71 |
| $g_s\ oscillation\ max$ | Maximum $g_s\ thermal$ reached as stomata re-opened at low *PPFD* | 0.20 | 0.07 | 0.08 | 0.53 | 0.67 |
| $(g_s\ oscillation\ max - g_s\ initial\ min)$ | Amplitude of oscillation of $g_s\ thermal$ at low *PPFD* | 0.15 | 0.05 | 0.05 | 0.43 | 0.49 |
| $t_{initial\ min}$ | Time for $g_s\ thermal$ to reach 110% of $g_s\ initial\ min$ | 5.2 | 1.6 | 1.8 | 13.4 | 0.68 |
| $g_s\ \Sigma\ shade$ | Area beneath the $g_s\ thermal$ curve following the *PPFD* decrease | 84 | 32 | 28 | 231 | 0.72 |
| $V_{initial}$ | Exponential rate of decline in $g_s\ thermal$ after the *PPFD* decrease | -0.89 | 0.27 | -2.23 | -0.08 | 0.67 |
| $V_{oscillation}$ | Linear rate of increase in $g_s\ thermal$ as stomata re-opened at low *PPFD* | 0.0052 | 0.0028 | 0.0006 | 0.022 | 0.24 |

### Genetic variation, heritability, and trait correlations

Variation among accessions was 3-fold, 10-fold, 6-fold, and 8-fold, respectively, for steady-state *g_s thermal_* at high *PPFD* (*g_s light_*), steady-state *g_s thermal_* at low *PPFD* (*g_s shade_*), the amplitude of decline in *g_s thermal_* from high to low *PPFD* (*g_s light_* - *g_s shade_*), and integrated *g_s thermal_* throughout the low-*PPFD* period (*g_s Σ shade_*, Table 1). For example, *g_s light_* in accession PI552851 was more than double that of accession PI267653 (Fig. 4A-B). After the decrease in *PPFD*, *g_s thermal_* declined at an exponential rate (*V_initial_*), then reached a minimum (*g_s initial min_*), and the time to reach this minimum was recorded (*t_initial min_*). Variation among accessions was 28-fold, 9-fold, and 7-fold, respectively, for *V_initial_*, *g_s initial min_*, and *t_initial min_*, respectively (Table 1). For example, the decline of *g_s thermal_* was faster in PI267653, which displayed a strongly negative *V_initial_* and low *t_initial min_* relative to slower accessions such as PI552851 (Fig. 4A-B). Stomatal re-opening often occurred after the initial decline causing a dampened oscillation in *g_s thermal_* during adjustment to low *PPFD*. The variation among accessions was 37-fold, 7-fold, and 9-fold, respectively, for the linear increase in *g_s thermal_* (*V_oscillation_*), *g_s thermal_* at peak stomatal re-opening (*g_s oscillation max_*) and the amplitude of oscillation (*g_s oscillation max_* - *g_s initial min_*). For example, the oscillation of *g_s thermal_* was more pronounced in NSL50717, which displayed high *V_oscillation_* and high oscillation amplitude (*g_s oscillation max_* - *g_s initial min_*), relative to PI660605, which showed no stomatal re-opening (Fig. 4C-D). Oscillation of *g_s thermal_* occurred on different timescales, e.g. more protracted in PI329646 than PI660630 (Fig. 4E-F). Variation in these traits among genotypes was reproducible across replicates (Fig. 4A-F).

**Figure 4:**
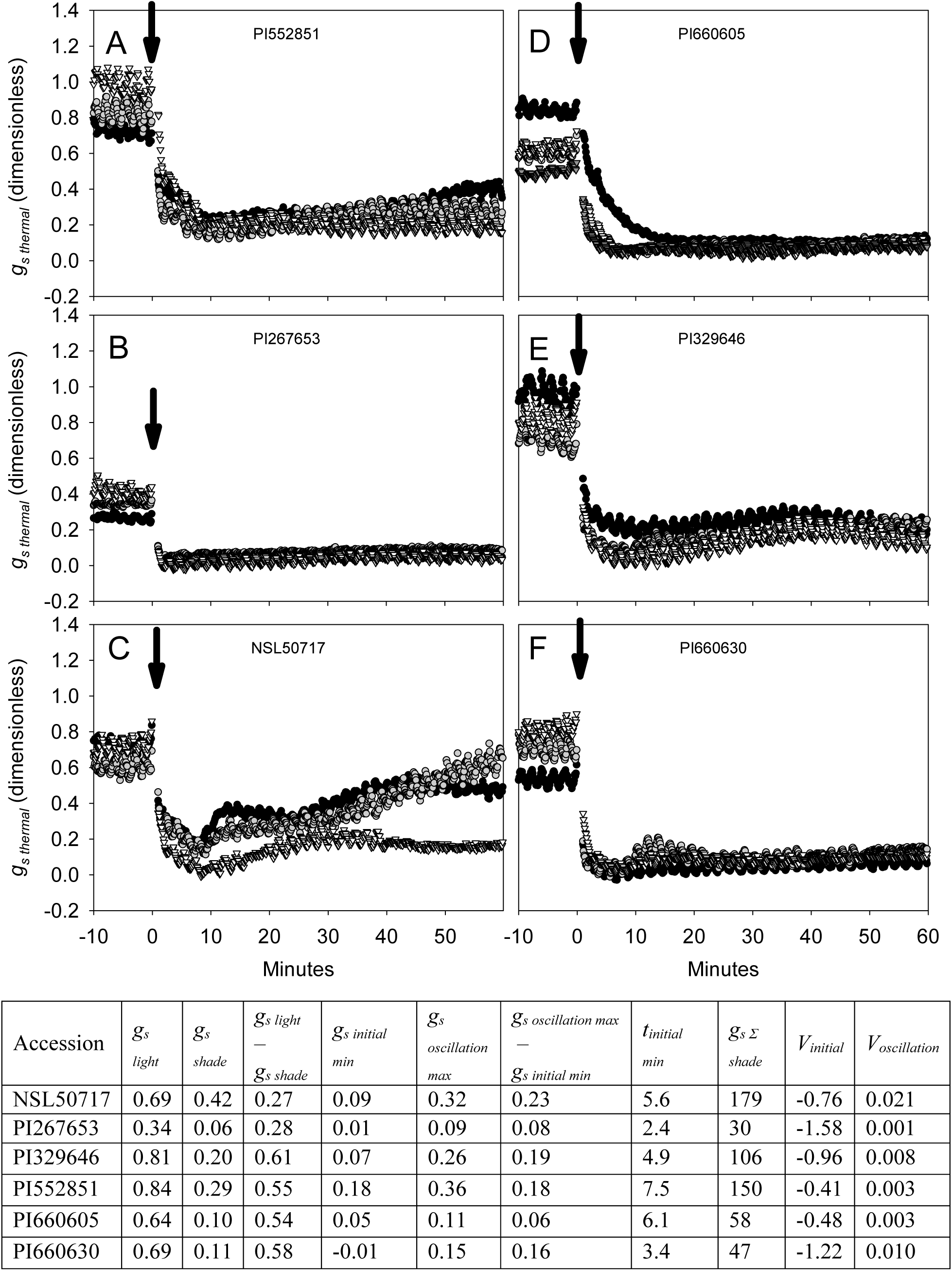
Representative responses of *g_s thermal_* to a 90% decrease in *PPFD* in six accessions, selected to highlight the diverse stomatal responses to decreasing *PPFD* measured in the sorghum diversity panel. Different symbols and shades of gray show distinct replicate plants for each accession. Each *g_s thermal_* response curve was measured on a single leaf, acclimated to *PPFD*=750 µmol m^-2^ s^-1^ for 40 minutes, then *PPFD* was reduced by 90% for 60 minutes at t=0, indicated by an arrow. Values in the table are the mean of stomatal conductance traits for each accession. *t_initial min_* is in minutes, all other traits are dimensionless.

Genomic heritability (*h^2^*) was highest (0.67-0.72) in traits describing *g_s thermal_* at low *PPFD*, including *g_s initial min_*, *g_s oscillation max_*, *g_s shade_*, and *g_s Σ shade_* (Table 1). Traits describing the speed of change in *g_s thermal_* after a decrease in *PPFD* (*V_initial_*, *t_initial min_*) also had moderately high *h^2^* (0.67-0.68). Traits describing the oscillation in *g_s thermal_* during stomatal re-opening at low *PPFD* (*V_oscillation_*, *g_s oscillation max_ - g_s initial min_*) had low to intermediate *h^2^* (0.24-0.49). *h^2^* was low (0.31) in traits describing *g_s thermal_* at high *PPFD* and the amplitude of overall decline in *g_s thermal_* from high to low *PPFD*: *g_s light_* and *g_s light_ - g_s shade_*, respectively.

All traits were correlated (*r* from −0.29 – 0.94) with one another (*p*<0.05, Fig. 5, Supplementary Fig. S1). In particular, accessions with high *g_s thermal_* at high *PPFD* also had greater overall *g_s thermal_* at low *PPFD* (positive correlation of *g_s light_* and *g_s Σ shade_*, *p*<0.0001, *r*=0.58, Fig. 6A), took longer to adjust to the decrease in *PPFD* (positive correlation of *g_s light_* and *t_initial min_*, *p*<0.0001, *r*=0.3; positive correlation of *g_s light_* and *V_initial_*, *p*<0.0001, *r*=0.34, Fig. 6B-C), and had more pronounced stomatal re-opening at low *PPFD* (positive correlation of *g_s light_* and *V_oscillation_*, *p*<0.0001, *r*=0.21, Fig. 6D).

**Figure 5:**
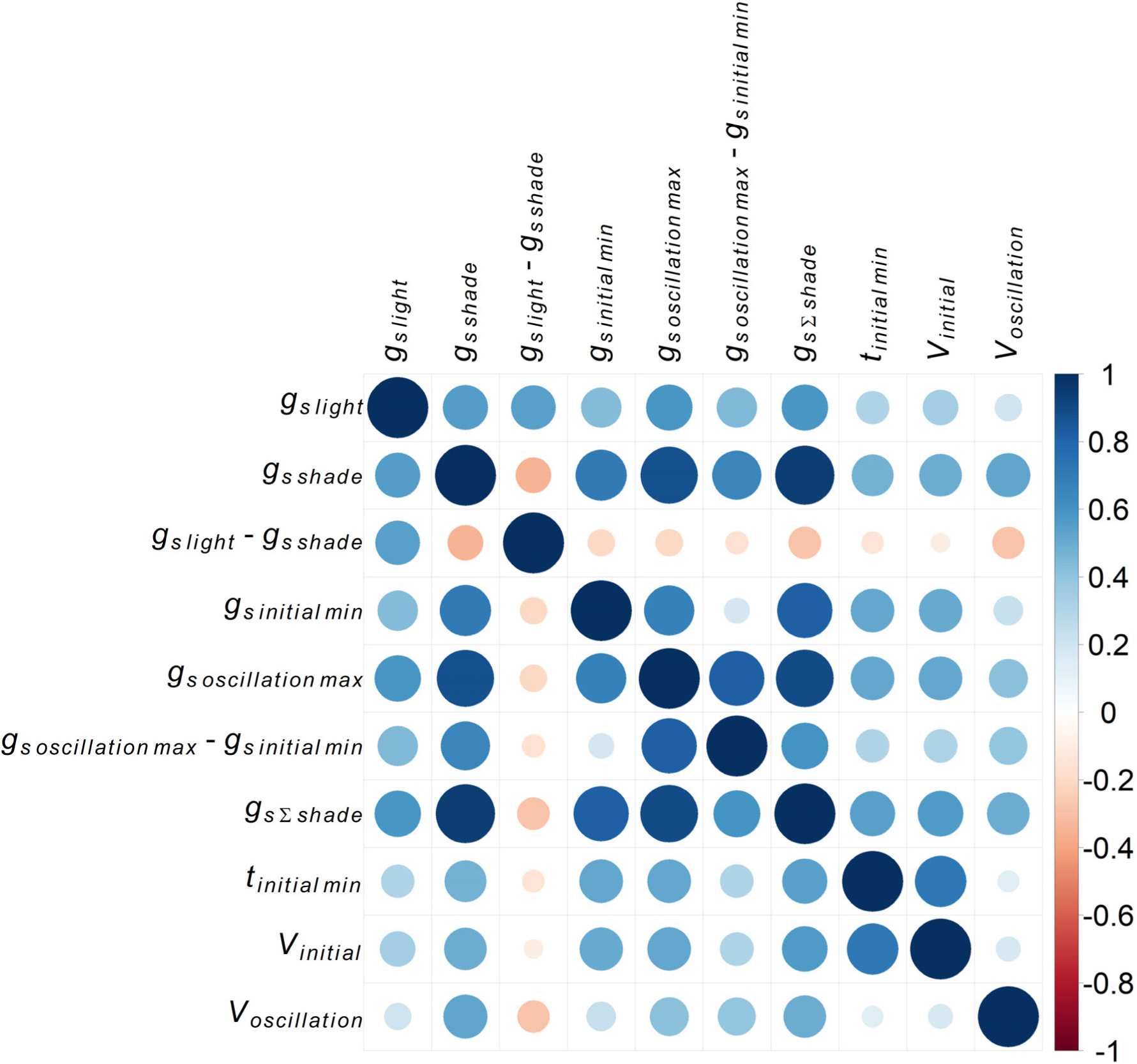
Pearson’s correlation coefficients (*r*) between BLUPs of stomatal conductance traits. The size of circles indicates the strength of correlation, while color indicates whether the pairwise relationship was negative (*r*<0, red) or positive (*r*>0, blue). All corresponding scatterplots are given in Supplementary Fig. S1.

**Figure 6:**
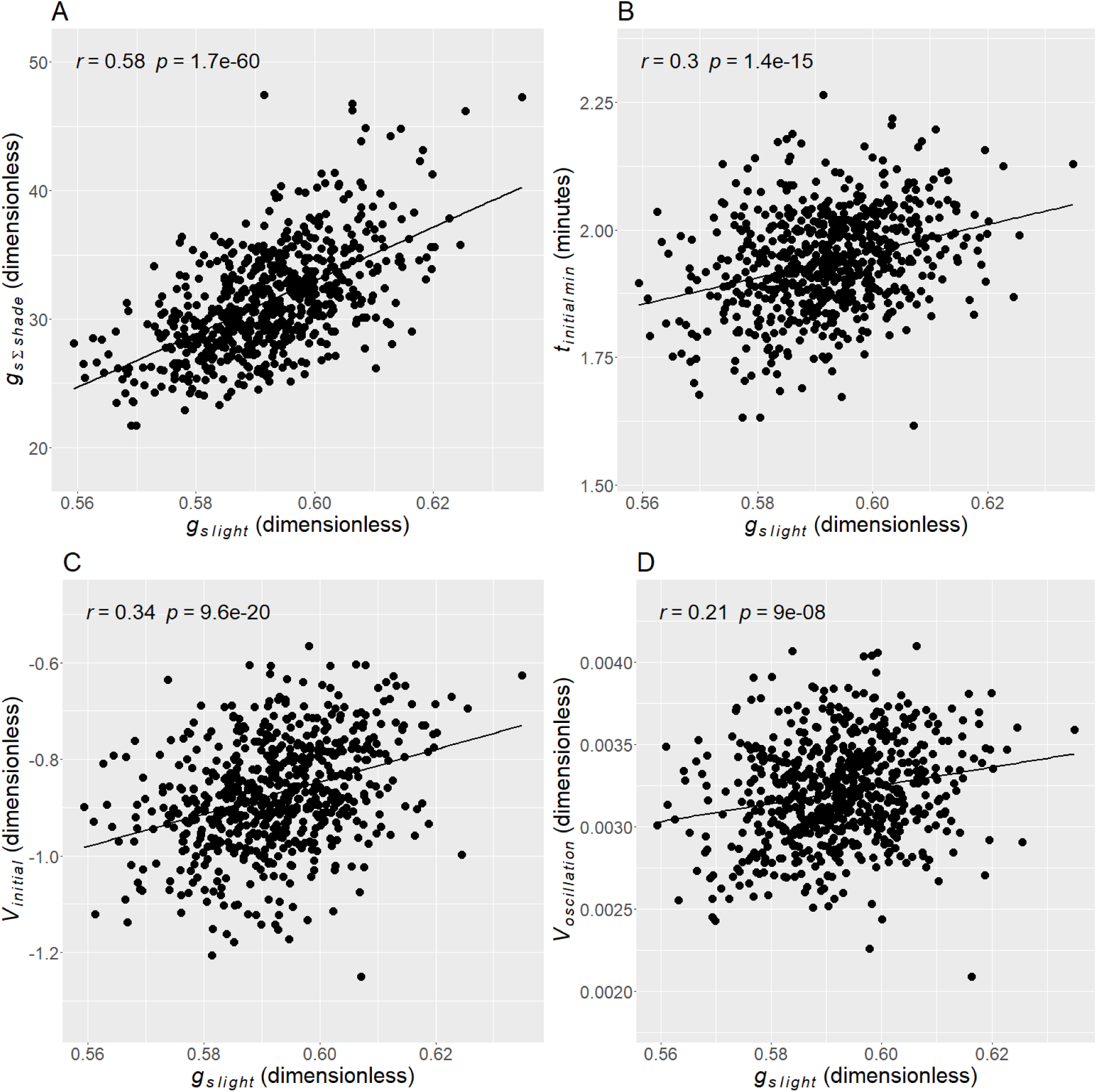
Correlation scatterplots between BLUPs for stomatal conductance traits. Pearson’s *r* and the associated *p*-value are also given.

### GWAS, TWAS, FCT and GO enrichment analysis

For each *g_s thermal_* trait, genes were initially identified as of potential interest if they were in linkage disequilibrium (LD) (Supplementary Table S1) with the top 0.1% strongest associated SNPs from GWAS (∼600 genes per trait; Supplementary Table S2); among the top 1% strongest associated genes from TWAS for leaf or shoot tissues (199 and 169 genes per trait, respectively; Supplementary Table S3); or among the top 1% strongest associated genes from FCT for leaf or shoot tissues (150 genes per trait; Supplementary Table S4). In addition to individual traits, multi-trait associations were performed with two trait groups: G1) traits describing the speed of change in *g_s thermal_* after a decrease in *PPFD* (*V_initial_*, *t_initial min_*), and G2) traits describing overall values of *g_s thermal_* (*g_s light_*, *g_s initial min_*, *g_s oscillation max_*, *g_s shade_*, *g_s Σ shade_*). The compilation of these results (Table S5) indicated that there was significant overlap in genes identified for different *g_s thermal_* traits, with 37% of genes being identified for two or more traits (Table 2).

**Table 2:** N. of genes overlapping the top results of GWAS, TWAS and/or FCT in multiple traits.

| **N. of overlapping traits** | **N. of genes** |
| --- | --- |
| 1 | 4575 |
| 2 | 1463 |
| 3 | 689 |
| 4 | 281 |
| 5 | 150 |
| 6 | 97 |
| 7 | 38 |
| 8 | 11 |
| 9 | 6 |
| 10 | 1 |

Follow-up analyses were also performed to identify a subset of “higher confidence” genes, i.e. genes for which there was evidence for an association of trait variation with DNA sequence variation (GWAS) and RNA transcript abundance (TWAS), or genes identified in tests from both of the two independent approaches to sampling developing leaf tissue i.e. from the 3^rd^ *leaf* or *shoot* section containing the growing point. Therefore, genes overlapping the top hits for multiple analyses/tissues, i.e. GWAS, TWAS leaf, TWAS shoot, FCT leaf, and/or FCT shoot (n. overlaps ≥2 in Supplementary Table S5), were further investigated (Supplementary Fig. S2-S12). Across all traits, 1548 candidate genes were identified consistently across two or more of the individual tests. Taking *V_initial_* as a representative trait, of the 1007 top hits, 180 were identified from at least two analyses/tissues (Fig. 7).

**Figure 7:**
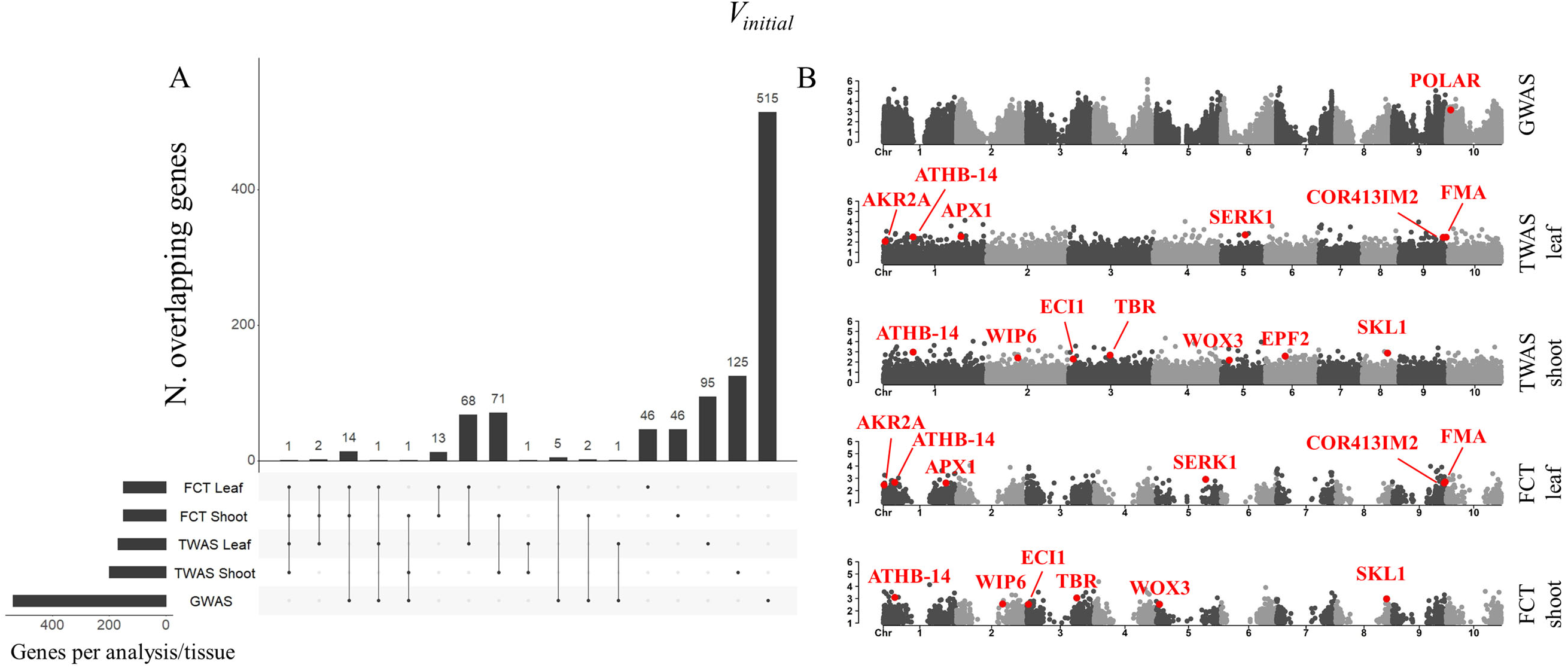
GWAS, TWAS and FCT results for *g_s light_*. A: Upset plot showing the number of overlapping genes between the top hits in GWAS, TWAS leaf, TWAS shoot, FCT leaf, and/or FCT shoot. B: Manhattan plots of GWAS, TWAS and FCT results. Top hits are highlighted if they correspond to orthologues of known Arabidopsis stomatal genes (FMA, POLAR and EPF2, Table 3). Top hits are also highlighted if they are among the highest confidence genes identified by GWAS, TWAS, FCT, and subsequent GO enrichment analysis. (Supplementary Table S7). Equivalent figures for all other traits are in Supplemental Figures S2-12.

GO enrichment analysis on Arabidopsis orthologs of the “higher confidence” genes identified 153 significantly enriched biological processes (FDR-adjusted *p*<0.05), nested within 34 broad categories (Supplementary Table S6). Among these, 22 categories of biological processes (e.g. *regulation of histone H3-K27 methylation*, *hydrogen peroxide metabolic process*, *cell cycle DNA replication*, *plant epidermis morphogenesis*, *lipid catabolic process* and *plant-type cell wall biogenesis*) were enriched by >2.5-fold (Fig. 8). A total of 239 unique genes contained within these GO categories were considered the strongest candidates to underlie variation in stomatal conductance traits studied here. A survey of the literature on their orthologs in Arabidopsis, maize and rice revealed a large proportion of genes (32 %) had functions related to stomatal opening/closing (24 genes), stomatal/epidermal cell development (35 genes), leaf/vasculature development (12 genes), or chlorophyll metabolism/photosynthesis (8 genes) (Supplementary Table S7).

**Figure 8:**
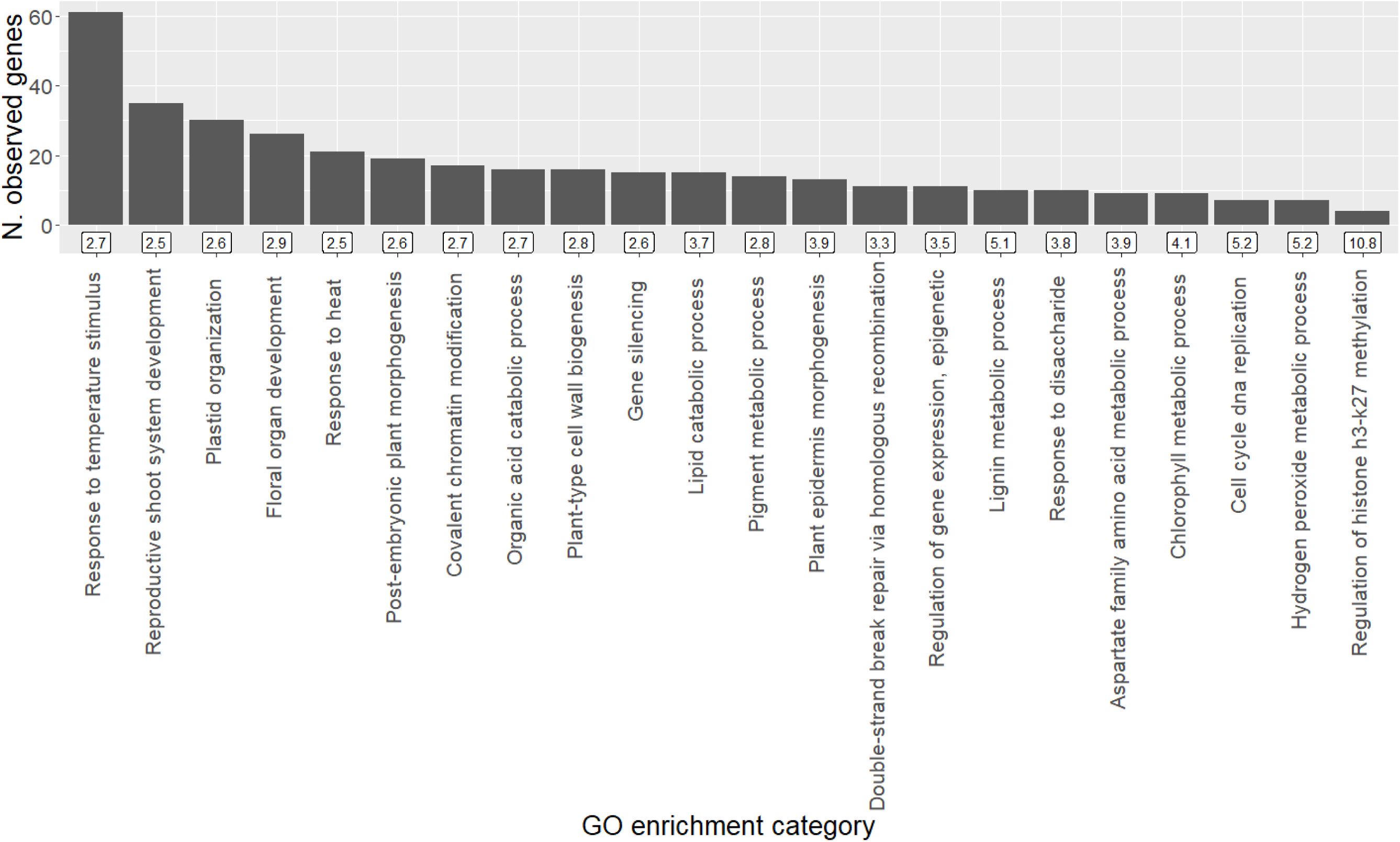
Results of GO enrichment analysis of higher confidence genes. Bars gives the number of genes included in each of the categories of GO biological processes that were significantly and >2.5-fold enriched. Fold-enrichment is shown in the label beneath each bar. Full GO enrichment analysis results are in Supplementary Table S6.

## Discussion

Stomatal responses to changes in *PPFD* influence WUE of plants in fluctuating light environments (Lawson and Blatt, 2014). This study successfully met the objectives of: (1) demonstrating how high-throughput thermal imaging can be used to rapidly phenotype variation in stomatal closure responses to PPFD across a diverse population of C_4_ plants; and (2) using that HTP data to perform an integrated GWAS/TWAS that identified a compelling set of candidate genes for further investigation of stomatal opening and closing in C_4_ species. Thermal measurements were validated against classical gas-exchange, and phenotypic correlations revealed relationships between steady-state (e.g. *g_s light_*) and dynamic (e.g. *V_initial_*) stomatal conductance traits. Results showed substantial, heritable variation in dynamic responses of stomata to a reduction in *PPFD*. This study presents important new information on sorghum, a model system with rapid stomatal movement compared to other species (McAusland et al., 2016), and addresses a major knowledge gap that exists for C_4_ species, despite their agricultural and ecological importance (Edwards et al., 2010; Leakey et al., 2019).

### Validation of *g_s thermal_* as a high-throughput proxy for *g_s_*

Although dynamic *g_s_* responses have been increasingly studied in the past few years (McAusland et al., 2016; Deans et al., 2019; Acevedo-Siaca et al., 2020; De Souza et al., 2020; Pignon et al., 2021), measurements have not yet been deployed at a scale amenable to association mapping (e.g., GWAS). Here, *g_s thermal_* was a useful high-throughput proxy for *g_s_*, enabling simultaneous measurement of 18 plants in a single imaging frame. Previous studies have reported near-perfect correlation between *g_s thermal_* and *g_s_* when it was examined globally across an entire dataset i.e. where *g_s_* varied by more than an order of magnitude as a result of combining data from multiple PPFDs, species and genotypes (Spearman’s rank-order ρ=0.96) (McAusland et al., 2013). In contrast, the present study focused on testing the correlations between estimates of individual traits measured by photosynthetic gas exchange versus thermal imaging at higher throughput. For example, thermal imaging was highly significant (p<0.001) in capturing variation in *g_s_* measured by gas exchange at a single PPFD (ρ ranging from 0.29 - 0.66, Fig. 3) and this thermal estimate of *g_s shade_* had a heritability of 0.70, making it suitable for association mapping. It is also notable that correlations between the two measurement approaches would have been weakened because *g_s thermal_* and *g_s_* were not measured simultaneously in the manner that McAusland et al. (2013) achieved so elegantly. For example, consistency between the two approaches to *g_s_* measurements was likely reduced by differences in measurement conditions and the immediate history of environmental conditions experienced by leaves as they moved from thermal measurements to the gas exchange chamber. Boundary layer conductance was likely lower while measuring *g_s thermal_* than *g_s_* due to the air mixing fan used in gas-exchange equipment (Grant et al., 2006). Light quality was equal parts red/blue/green for *g_s thermal_* vs. 90% red 10% blue for *g_s_*. This likely affected stomatal opening, which is induced by blue, and to a lesser extent, red light (Assmann and Shimazaki, 1999; Shimazaki et al., 2007; Lawson et al., 2011; Assmann and Jegla, 2016). For these reasons, the significant correlations between all traits estimated from *g_s thermal_* and their counterparts estimated from gas exchange measurements were considered a validation of the methods used.

### Sorghum shows varied, heritable stomatal responses to a decrease in *PPFD*

Within-species diversity in stomatal light responses has been documented for C_3_ species including Arabidopsis (Takahashi et al., 2015), poplar (Durand et al., 2019), rice (Acevedo-Siaca et al., 2020), cassava (De Souza et al., 2020) and soybean (Soleh et al., 2016). Expanding the scale of investigation to 659 accessions of sorghum revealed substantial variation within the C_4_ model species, despite the fact that C_4_ species generally have lower *g_s_* than C_3_ species (Taylor et al., 2010). The range of *t_initial min_* shown here (1.8-13.4 minutes, Table 1) overlapped with similar measurements in sorghum (*∼*2-8 minutes) (McAusland et al., 2016; Pignon et al., 2021), the closely related C_4_ grasses miscanthus and maize (*∼*8 minutes), the C_3_ grass rice (*∼*10 minutes) (McAusland et al., 2016), and the semi-aquatic rhizomatous fern *Marsilea drummondii* A. Braun (*∼*9 minutes) (Deans et al., 2019). Some C_3_ dicots such as Arabidopsis and sunflower (*∼*18 minutes) were roughly comparable to the slowest accession shown here (*t_initial min_* =13.4 minutes), while many other species appeared considerably slower (*∼*30 minutes) (McAusland et al., 2016; Deans et al., 2019). The faster stomata of sorghum might be related to the unique structure of graminaceous stomata, composed of dumbbell-shaped guard cells flanked by subsidiary cells, which have been linked to rapid movement relative to other forms (Franks and Farquhar, 2007; Lawson et al., 2011; Serna, 2011; McAusland et al., 2016; Lawson and Vialet-Chabrand, 2019). Stomata of C_4_ plants tend to be smaller and more sensitive to environmental change than their C_3_ counterparts (Lawson et al., 2011; McAusland et al., 2016).

### Implications of natural diversity in stomatal responses to decreasing *PPFD*

Compared to steady-state *g_s_* under high *PPFD,* traits describing the dynamic change in *g_s thermal_* after a decrease in *PPFD* had greater variability and *h^2^*, making them more tractable targets for association studies (Table 1). Additionally, variation in traits describing the speed of change in *g_s thermal_* after a decrease in *PPFD* (*V_initial_*, *t_initial min_*) might be leveraged to accelerate stomatal responses even in species with “fast” stomata such as sorghum, potentially improving coordination of *g_s_* with photosynthetic carbon assimilation (*A*) and resulting in improved *iWUE* (Lawson and Blatt, 2014).

The relationships among traits observed across the genetic variation surveyed here are consistent with biophysical trade-offs driven by structure-functional relationships, as well as selection for trait combinations that favor carbon gain versus water savings to differing degrees in different environments. For example, the finding that accessions with greater *g_s light_* took longer to adjust to a decrease in *PPFD* (Fig. 6B-C) is consistent with greater *g_s light_* being associated with larger stomata and longer times for pores to close in a subset of the lines studied here (Pignon et al., 2021) and in tests of diverse species (McAusland et al., 2016). However, it is worth noting that variation in stomatal opening/closing is also associated with guard cell physiology, including ion transport processes (Lawson and Blatt 2014). Adaptation to different environments may also contribute to the observed trait correlations, with high *g_s light_* and slow stomatal closure working in concert to favor *A* and rapid growth in environments where water is not limiting. In contrast, low *g_s light_* and rapid stomatal closure combine to prioritize water-use efficiency and conservative but sustained growth in water-limited environments (Vico et al., 2011). Since more rapid stomatal closure after a decrease in PPFD would increase iWUE, there is significant interest in identifying more genes underpinning structural and functional components of stomatal movements, as well as their interactions with steady-state gas exchange and leaf development and physiology more broadly (Lawson and Blatt 2014).

### GWAS/TWAS identifies genes enriched in stomatal, leaf developmental and photosynthetic functions

Gene candidates putatively associated with genetic variation in stomatal closure in sorghum were identified using GWAS and TWAS integrated with FCT, followed by GO enrichment analysis. This approach has identified known causal variants more efficiently than GWAS and TWAS alone (Kremling et al. 2019), while also increasing the consistency in results observed when testing was repeated across different conditions (Ferguson et al. 2020). The present study reinforced these prior reports, with an order of magnitude more genes being consistently identified by FCT versus TWAS across the two independent tissue sampling strategies used (Table S5). GO enrichment analysis of the Arabidopsis orthologues of these genes revealed 22 GO biological processes that were significantly and >2.5-fold enriched (Supplementary Table S6, Fig. 8). The 239 genes belonging to these 22 categories were selected as the greatest confidence candidate genes (Fig. 8; Supplementary Table S7). A large proportion (32%) of these genes have orthologs in Arabidopsis, maize or rice that are already implicated in regulating traits related to stomata or WUE. While it is unlikely that such enrichment would occur by random chance, the function of the genes identified here will require confirmation by follow-up reverse genetic studies of transgenic or mutant plants.

Twenty three orthologs of genes implicated in signaling, metabolism or transporters in guard cells belong to enriched GO terms including *lipid catabolic process*, *hydrogen peroxide metabolic process*, *response to disaccharide* and *response to heat* (Supplementary Table S7). For example, loss of ascorbate peroxidase 1 (APX1) and the Respiratory burst oxidase homolog protein F (RBOHF) influence redox oxygen species (ROS) to alter stomatal responses to light, [CO_2_] or abscisic acid (ABA; Pnueli et al., 2003; Chater et al., 2015; Sierla et al., 2016). Meanwhile, the MYB60 transcription factor is required for light induced opening of stomata in Arabidopsis. It is expressed exclusively in guard cells, with expression increasing and decreasing in accordance with conditions that promote stomatal opening and closing, respectively (Cominelli et al., 2005). Phospholipase Dα1 (PLDα1) and its lipid product phosphatidic acid impact ABA-induction of ROS production and stomatal closure (Zhang et al., 2009). Mutants of ABC transporter G family member 40 (ABCG40) shut more slowly in response to ABA (Kang et al. 2010), while its sorghum ortholog was variously associated with 5 different *g_s_* thermal traits in GWAS, TWAS and FCT tests.

Recently, lipid metabolism of guard cells was discovered to be important as an energy source for light-induced stomatal opening (McLachlan et al., 2016). Five sorghum genes associated with variation in *g_s thermal_* traits were orthologs of genes involved in triacyl glyceride mobilization and expressed in guard cells of Arabidopsis (Enoyl-CoA delta isomerase 1 and 3, ECI1 and ECI3; peroxisomal fatty acid beta-oxidation multifunctional protein, MFP2; Acyl-coenzyme A oxidase 2 and 4, ACX2 and ACX4; McLachlan et al., 2016). Notably, Wrinkled1, a transcription factor that regulates metabolic genes in a manner that promotes carbon allocation to fatty acid synthesis, was also identified (Cernac and Benning 2004). Therefore, a significant proportion of the highest confidence candidate genes identified by the integrated GWAS/TWAS are plausibly involved in signaling, metabolism and transport functions in guard cells. Further study will be needed to determine if the candidate genes identified here play specific roles in stomatal closure, or if the associations observed in this study are partly a product of the strong correlations between rates of stomatal opening and closing (Lawson and Blatt 2014). Current understanding of lipid metabolism guard cells would seem to suggest the latter option is more likely, but this area of study is still relatively nascent.

The functions of orthologs of the highest confidence sorghum candidate genes are also consistent with the importance of structure-function relationships to the speed of stomatal opening and closing. Thirty five orthologs of genes implicated in stomatal or epidermal cell patterning belong to enriched GO terms including *plant epidermis morphogenesis*, *cell cycle DNA replication*, *plant-type cell wall biogenesis*, and *plastid organization* (Supplementary Table S7). Arabidopsis genes known to impact stomatal development or patterning which had sorghum orthologs found in the highest confidence candidates for *g_s thermal_* traits included: the Mitogen-activated protein kinase kinase kinase YODA (Bergmann et al., 2004), the FAMA bHLH-type transcription factor (Ohashi-Ito and Bergmann 2006), the cyclin-dependent kinase CDKB1;1 (Boudolf et al., 2004), the phytochrome interacting factor 1 (PIF1, Klermund et al., 2016), somatic embryogenic receptor kinase 1 (SERK1, Meng et al. 2015), DNA-directed RNA polymerase II subunit 2 (Chen et al., 2016), the chromatin regulator Enhanced Downy Mildew 2 (EDM2, Wang et al., 2016), Protein Phosphatase 2A (Bian et al., 2020), GATA, NITRATE-INDUCIBLE, CARBON METABOLISM-INVOLVED (GNC, Klermund et al., 2016), the extra-large GTP-binding protein (XLG3, Chakravorty et al., 2015), and the ARF guanine-nucleotide exchange factor GNOM (Le et al., 2014). Focusing on grasses, SOBIC.010G277300 shares 96 % predicted protein sequence homology with GRMZM2G057000 in maize. The *nana plant2* (na2) mutant of this gene displays alterations in brassinosteroid synthesis and the morphology of stomatal complexes (Best et al., 2016). Similarly, SOBIC.004G116400 shares 74 % predicted protein sequence homology with Os02g15950 (ERECT PANICLE 3, EP3) in rice. Loss of function mutants of EP3 have smaller stomata, which appeared to drive reductions in *g_s_* and *A* (Yu et al., 2015). Given the substantial evidence for links between gas exchange, stomatal complex size and stomatal density (Lawson and Blatt 2014; Xie et al., 2020), there is potential for variations in the sequence and expression of these genes to drive variation in the *g_s thermal_* traits measured in this study. A number of other genes identified by the integrated GWAS/TWAS have been implicated in the development of epidermal cells in general (Supplementary Table S7). Cross talk between development pathways for stomata and other types of epidermal cells (Kim and Dolan 2011, Raissig et al., 2016) creates the opportunity for such genes to influence *g_s_* and its response to PPFD.

Finally, a smaller number of sorghum genes identified in this study have orthologs known to influence overall leaf development/vasculature (9 genes) or chlorophyll/photosynthesis (7 genes) (Supplementary Table S7). Leaf vasculature determines the hydraulic capacity of the leaf to deliver water that eventually diffuses out of the leaf as vapor. Consequently, strong traits associations between leaf hydraulics and stomata have been described in a range of contexts (Sack et al., 2003; Bartlett et al., 2016). In that vein, it is plausible that genes known to alter vascular development via effects on polyamine metabolism (5’-methylthioadenosine nucleosidase, MTN1, Waduwara-Jayabahu et al., 2012), glucuronoxylan synthesis (Beta-1,4-xylosyltransferase, IRX9, Pena et al., 2007) and transcriptional regulation (DEFECTIVELY ORGANIZED TRIBUTARIES 5, DOT5, Petricka et al., 2008) might be associated with variation in *g_s_* thermal traits. Similarly, identification of genes involved in photosynthesis may reflect the tight linkage between *A* and *g_s_*, which is observed in many plant species and is particularly strong in sorghum (Leakey et al., 2019). QTL for traits related to *A* and *g_s_* often overlap, including in sorghum (Ortiz et al., 2017). When compared to other species, sorghum shows exceptional coordination between *g_s_* and *A* following decreases in *PPFD*, driven by rapid responses in *g_s_* (McAusland et al., 2016). Among diverse sorghum accessions, there is significant covariation between the responses of *A* and *g_s_* following decreases in *PPFD*, i.e. accessions with more rapid declines in *A* also have more rapid declines in *g_s_*, and vice-versa (Pignon et al. 2021). Most notable was the identification of the sorghum Rubisco Activase (RCA) by GWAS, TWAS and FCT across 8 different *g_s thermal_* traits (Supplementary Table S7). RCA encodes an enzyme that plays a key role in activating Rubisco to perform the key step in photosynthetic CO_2_ assimilation (Portis 2003), and which is known to limit the rate of photosynthetic induction after an increase in PPFD (Pearcy 1990). Along with a number of the results described above, this opens the possibility that phenotyping only stomatal closure may have facilitated identification of associations between genotype or gene expression and both stomatal opening and closing, as a result of the two aspects of stomatal movement being so tightly linked. If functional validation of candidate genes supports that notion, then considerable time can be saved when collecting phenotypic data.

## Conclusion

This study demonstrates how high-throughput phenotyping by thermal imaging can be used to assess genetic variation in stomatal closure after a decrease in PPFD at a scale suitable for association mapping. Integrated GWAS/TWAS and FCT was then applied to identify a set of candidate genes, which were enriched in orthologs of Arabidopsis, maize and rice genes involved in stomatal opening/closing, epidermal patterning, leaf development and photosynthesis. This is important proof of concept for methods to break the phenotyping bottleneck for a trait that is important to plant productivity and sustainability but has until now been intractable as a target for study by quantitative genetics. The method described here could also be applied to other species from a variety of plant functional types or grown in different environments to assess G×E of stomatal traits. In addition, the study provides new knowledge of trait variation and underlying candidate genes in an important C_4_ model crop, which is notable for the speed of its stomatal opening/closing. This lays the foundation for future studies to establish gene function and potentially improve crop performance.

## Materials and methods

### Physiology measurements

#### Plant material and growing conditions

A random subset of 659 accessions was selected from the biomass sorghum diversity panel at the University of Illinois at Urbana-Champaign, as previously described (Valluru et al., 2019). Accession names are provided in Supplementary Table S8. Plants were grown from seed in flats (3×6 sets of 281 mL inserts) containing a peat/bark/perlite-based growing medium (Metro-Mix 900; Sun Gro Horticulture, Agawam, MA, USA) and supplemented with 1 mL slow release 13-13-13 fertilizer (Osmocote Classic, Everris NA, Inc., Dublin, OH, USA). One accession, PI147837, was included in each flat to identify spatial and temporal variation in measurements. Three seeds/insert were planted, then thinned to 1 seedling/insert. Flats were watered regularly to field capacity and grown in a greenhouse maintained at 27 °C day/25 °C night, with supplemental lighting to ensure minimum light intensity of 90 W m^-2^ during a 13 h day. n=3-4 plants were assessed per accession.

#### Experimental conditions and leaf temperature measurement

Once the fourth leaf had fully expanded, as evidenced by ligule emergence, plants were transferred to a growth cabinet overnight (Model PCG20, Conviron, Winnipeg, MB R3H 0R9, Canada). Cabinets were maintained at 14 h/10 h day/night cycle under 1200 μmol photons m^−2^ s^-1^ PAR, 30 °C daytime/25 °C nighttime temperature, and 75% RH. On the day of measurement, the fourth leaf of each plant was laid flat across a frame to standardize leaf angle and incident light interception (Supplementary Fig. S13). This presented a 4 cm length of the mid-leaf for measurement. Leaves were not detached from plants. Dry and wet reference materials were prepared as in McAusland et al. (2013) to correct for the effects of net isothermal radiation and VPD, respectively (Guilioni et al., 2008). A thin coating of petroleum jelly was applied over 1 cm of abaxial and adaxial sides of leaves, providing a dry reference unique to each leaf. Two sections of filter paper, moistened by a water reservoir, were used as a wet reference.

Flats of 18 plants were transferred to a second growth cabinet with conditions identical to the first cabinet, except that light was provided by a 20 x 20 cm LED panel providing equal-parts blue, red and green light, with a combined incident photon flux of 750 µmol m^-2^ s^-1^ at the leaf level (LED Light Source SL 3500, Photon Systems Instruments, Brno, Czech Republic). An infrared camera (Thermo Gear Model G100, Nippon Avionics CO., Ltd., Tokyo, Japan) was placed 0.5 m above the leaves without obstructing the light source. Incident photon flux was maintained at 750 µmol m^-2^ s^-1^ for 40 minutes, and then reduced by 90% for an additional 60 minutes. Images were recorded every 6 seconds, with emissivity=0.95 (Jones et al., 2002). The cabinet’s PAR sensor was used to evaluate spatial heterogeneity of incident photon flux, which was contained to ± 6% variation across the measured area.

#### Image analysis and stomatal conductance estimation

Analysis of thermal images was performed in ImageJ (ImageJ1.51j8, NIH, USA).

Sections of leaf and reference materials of each image were hand-selected to derive profiles of temperature vs. experimental time. Leaf and reference temperatures were used to calculate *g_s thermal_*:

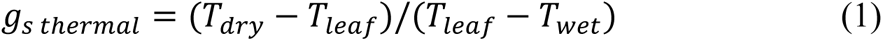

where *T_leaf_*, *T_dry_*, and *T_wet_* are temperatures of the leaf, dry and wet references, respectively. *g_s thermal_* is theoretically proportional to *g_s_* given constant environmental conditions (Jones, 1999; Jones et al., 2002; Grant et al., 2006; Guilioni et al., 2008; McAusland et al., 2013). Air RH and temperature were controlled by the growth cabinet and assumed constant across all leaf and reference surfaces. Since the cabinet was designed to deliver a uniform airflow, and replicate plantings were randomly positioned to avoid systematic spatial variation, boundary layer conductance was also assumed constant. Differences in *g_s thermal_* between leaves and over time were therefore attributed to *g_s_*.

### Analysis of *g_s thermal_* profiles

Several traits were derived from profiles of *g_s thermal_* vs. experimental time: *g_s light_*, *g_s shade_*, *g_s Σ shade_*, *g_s initial min_*, *g_s oscillation max_*, *t_initial min_*, *V_initial_*, and *V_oscillation_*. A graphical description of these traits is given in Fig. 1. After the decrease in *PPFD*, *g_s thermal_* declined as stomata closed, often followed by an oscillation in *g_s thermal_* as stomata re-opened and then closed again. *g_s light_* and *g_s shade_* were the average of *g_s thermal_* from t=-5 to 0, and from t=52 to 60 minutes, respectively. These gave steady-state *g_s thermal_* at *PPFD* of 750 and 75 μmol m^-2^ s^-1^, respectively. *g_s Σ shade_* was the area beneath the curve following the *PPFD* change, i.e. from t=0 to 60 minutes. *g_s initial min_* was the minimum of *g_s thermal_* reached immediately after the decrease in *PPFD*. *g_s oscillation max_* was the maximum of *g_s thermal_* reached during the stomatal re-opening phase. The time at which *g_s thermal_* reached 110% of *g_s initial min_* was recorded as *t_initial min_*. The amplitude of the overall change in *g_s thermal_* from high to low *PPFD* was (*g_s light_* - *g_s shade_*), and the amplitude of oscillation in *g_s thermal_* at low *PPFD* was (*g_s oscillation max_* - *g_s initial min_*).

*V_initial_* was derived from non-linear regression (PROC NLIN, SAS v9.4; SAS Institute, Cary, NC, USA) as the exponential rate of decline of *g_s thermal_* from t= −0.1 minutes to t= *t_initial min_*:

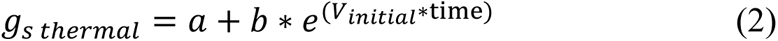

here *b* and *a* give estimates of *g_s thermal_* at t= −0.1 minutes and t=*t_initial min_*, respectively, and a more negative *V_initial_* indicates a more rapid decline in *g_s thermal_*. *V_oscillation_* was derived from linear regression (PROC GLM, SAS v9.4) as the linear slope of *g_s thermal_* vs. time during stomatal re-opening at low *PPFD*. A more positive *V_oscillation_* indicates a more rapid stomatal re-opening.

#### Validation of g_s thermal_ with gas-exchange measurements

Validation of *g_s thermal_* as a proxy for *g_s_* was obtained on a subset of 64 plants. After *g_s thermal_* measurements were completed, plants were placed back in the first growth cabinet. The leaf section previously used for *g_s thermal_* measurements was placed in the cuvette of a portable gas-exchange system incorporating infra-red CO_2_ and water vapor analyzers (LI-COR 6400; LI-COR, Inc., Lincoln, NE USA). Incident *PPFD* was set to 750 μmol m^-2^ s^-1^, [CO_2_] to 400 ppm, and leaf-to-air water vapor pressure deficit maintained <2 kPa. *PPFD* was 10% blue and 90% red light provided by integrated red and blue LEDs. The *g_s thermal_* measurement protocol was replicated, i.e. initial *PPFD* was maintained for 40 minutes, then reduced by 90% for an additional 60 minutes, with *g*_s_ logged every 5 seconds (von Caemmerer and Farquhar, 1981). Pearson’s correlation (*r*) at *p*=0.05 threshold, along with Spearman’s rank-order correlation (ρ) were tested between equivalent stomatal conductance traits derived from *g_s thermal_* and *g_s_* using R 3.6.1 (R Core Team, 2017).

When comparing *g_s thermal_* measurements to this validation data, an anomalous spike in *g_s thermal_*, reaching up to twice the steady-state high-*PPFD g_s thermal_*, was consistently recorded from t=0 to 0.9 minutes (Fig. 1). This was likely due to the different radiative properties of the white wet reference and the green leaf and dry reference. Therefore, *g_s thermal_* measurements from t=0 to 0.9 minutes were discarded.

#### Model development for best linear unbiased predictors (BLUPs)

A linear mixed model was used to account for spatial and temporal variation using the ASReml-R package (Butler et al., 2009). The best linear unbiased predictors (BLUPs) were obtained for all accessions and traits and were used for subsequent analysis (Supplementary Table S8). The most appropriate model for each trait was chosen in two steps. First, fixed effects with a Wald statistics *p*-value > 0.05 were excluded from the model. Subsequently, the Akaike information criterion (AIC) was used to select random effects variables and residual variance-covariance structures for each trait. The full model was:

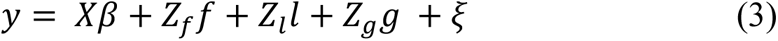

where *y* is the vector of phenotypes, *β* is a vector of fixed effects including the intercept, a blocking effect, and a cubic smoothing splines terms for leaf position and time of measurement, with design matrix *X*. Here the block term refers to three discrete periods throughout the experiment where the LED light had to be repaired and repositioned within the measurement cabinet. The vector *f* is the vector of random effects of “flat” within block with 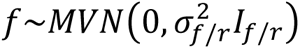 and design matrix *Z_f_* Here the term “flat” refers to a sequential flat number to account for temporal variation between measured flats of plants. The vector *l* is the vector of random effects of the interaction between row leaf position and column leaf position with 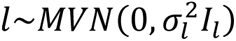 and design matrix *Z_l_.* The vector *g* is the vector of random genotypic effects of accessions with 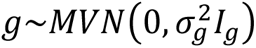 and design matrix *Z_g_* and *ξ* is the vector of residuals with distribution 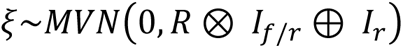. The matrix 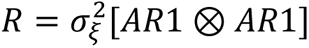 represents the Kronecker product of first-order autoregressive processes across row and column plant positioning within a flat, respectively, and 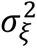 is the spatial residual variance. The matrices *I_f/r_*, *I_g_*, *I_l_* and *I_r_* are the identity matrices of the same dimensions as “flat” within block, genotypic effects, row leaf position and column leaf position interaction effect, and block, respectively. Outliers were removed following method 2 of (Bernal-Vasquez et al., 2016), and the Box-Cox power transformation was used on traits with non-normal residuals.

BLUPs were obtained for all accessions and each trait and added to the grand mean for GWAS and TWAS. Genetic and residual variances estimated from the null GWAS model were used to calculate genomic heritability (*h^2^*) as the ratio of genetic variance over phenotypic variance (de los Campos et al., 2015). Phenotypic correlations between all traits were tested using Pearson’s correlation (*r*) at *p*=0.05 threshold using cor.mtest() function in package corrplot (Wei and Simko, 2017).

### Genomic data collection for GWAS and TWAS

#### Genotypic data

Genotyping was performed as previously reported (dos Santos et al., 2020). Briefly, DNA from dark-grown etiolated seedling tissue was extracted and placed in 96-well plates following CTAB protocol (Doyle and Doyle, 1987). The genotyping was done using two pairs of restriction enzymes, PstI-HF/HinP1I and PstI-HF/BfaI (New England Biolabs, Ipswich, MA, USA) with the genotyping-by-sequencing (GBS) protocol (Elshire et al., 2011; Morris et al., 2013). Tag alignment was done with Bowtie2 (Langmead and Salzberg, 2012) using the *Sorghum bicolor* genome v3.1 (www.phytozome.jgi.doe.gov). SNPs were identified using the TASSEL3 GBS pipeline (Glaubitz et al., 2014). Reads that did not perfectly match a barcode and restriction site were discarded. After barcode trimming, all unique 64 bp sequences present >9 times in the dataset and that mapped uniquely to the sorghum genome were selected as “master tags.” These were compared to tags in each individual at each genomic address to identify SNPs. SNPs with >95% missing data or minor allele frequency (MAF)<5% were discarded.

A HapMap of 239 whole-genome-resequenced sorghum accessions containing 5.5M biallelic phased SNPs with MAF>0.01 (Valluru et al., 2019), was used as a reference panel to impute the GBS data. GBS markers were filtered to only consider markers that were also present in the HapMap. Beagle 4.1 (Browning and Browning, 2016) was used under GT mode with Ne set to 150,000, window=60,000 SNPs, and overlap=4,000 SNPs. After imputation, markers with AR2 < 0.3 were removed, resulting in 2,457,023 SNPs. LD pruning using PLINK (Chang et al., 2015) eliminated markers in high LD (*r*^2^ > 0.9) within 50kb windows. The final dataset consisted of 422,897 SNPs.

#### Gene expression data

186 sorghum accessions were grown for 3’RNAseq measurement. Environmental conditions were: 12 h/12 h day/night cycle under 500 µmol m^-2^ s^-1^ PAR, 25 °C daytime/23 °C nighttime temperature, and 75% RH. 2 cm of shoot and leaf tissues were sampled at 3^rd^ leaf stage. Samples were processed according to (Kremling et al., 2019). Briefly, RNA was extracted using TRIzol (Invitrogen) with Direct-zol columns (Zymo Research), and 3′ RNA-seq libraries were prepared robotically from 500 ng total RNA in 96-well plates on an NXp liquid handler (Beckman Coulter) using QuantSeq FWD kits (Lexogen) according to the manufacturer’s instructions. Libraries were pooled to 96-plex and sequenced with 90 nucleotide single-end reads using Illumina TruSeq primers on an Illumina NextSeq 500 with v2 chemistry at the Cornell University Sequencing facility.

The first 12 bp and Illumina Truseq adapter remnants were removed from each read using Trimmomatic version 0.32, following kit marker instructions. The splice-aware STAR aligner v.2.4.2a was used to align reads against the sorghum v3.1.1 reference genome annotations, allowing a read to map in at most 10 locations (-outFilterMultimapNmax 10) with at most 4% mismatches (–outFilterMismatchNoverLmax 0.04), while filtering out non-canonical intron motifs (–outFilterIntronMotifs RemoveNoncanonicalUnannotated). Default settings from STAR v.2.4.2a aligner were used to obtain gene-level counts (--quantModel GeneCounts) from the resulting BAM files.

### GWAS, TWAS and FCT

Traits went through an additional normal quantile transformation, then single and multi-trait associations were performed as in (Zhou and Stephens, 2014). Two groups of multi-trait models were considered: G1) traits describing the speed of change in *g_s thermal_* after a decrease in *PPFD* (*V_initial_*, *t_initial min_*), and G2) traits describing overall values of *g_s thermal_* (*g_s light_*, *g_s initial min_*, *g_s oscillation max_*, *g_s shade_*, *g_s Σ shade_*). Both groups of traits went through a step of multivariate outlier removal (Filzmoser et al., 2005) performed before running GWAS and TWAS.

GEMMA (Zhou and Stephens, 2012) was used for single and multivariate GWAS (Zhou and Stephens, 2014). Population structure was accounted for by using principal components (PCs) as fixed effects. Based on the Scree plot, 4 PCs obtained from PLINK (Chang et al., 2015) using the full SNP dataset (i.e. not LD-pruned) were included in all models. Relatedness was controlled for by a kinship matrix obtained from TASSEL 5 (Bradbury et al., 2007) using the default method (Endelman and Jannink, 2012).

TWAS was tested in developing leaf and shoot tissues with genes expressed in at least half of tested plants. Analyses were implemented in R 3.3.3 (R Core Team, 2017) with the *lm* function used for single-trait TWAS and the MANOVA function for multi-trait TWAS. Similar to (Kremling et al., 2019), 29 Peer factors (Stegle et al., 2010) and five multidimensional scaling factors were used as covariates.

An ensemble approach combining GWAS and TWAS results was performed using the Fisher’s combined test (FCT) (Kremling et al., 2019). Briefly, to integrate both the results from GWAS and TWAS, each SNP in the top 10% of GWAS analysis was assigned to the nearest gene. The *p*-values of genes not tested in the TWAS (genes expressed in less than half of tested plants) were set to one. The GWAS and TWAS *p*-values for each gene were combined using Fisher’s combined test in *metap* package in R, producing Fisher’s combined *p*-values.

#### Candidate gene analysis

Results of GWAS, TWAS and FCT were used to identify potential candidate genes driving variation in traits. A threshold set at the 0.1% lowest *p*-values was used to identify candidates for each SNP-trait association, i.e., 423 marker associations per trait. This threshold was chosen to focus the analysis on a minimum number of large-effect variants and to limit the number of false positives. PLINK (Chang et al., 2015) was used to calculate LD blocks with option --blocks and a window of 200 kb and default values for D-prime’s confidence interval (0.7;0.98) (Supplementary Table S1). Genes within these LD blocks were compiled from the Phytozome database for Sorghum bicolor v3.1.1 (Goodstein et al., 2012). Similarly, the top 1% most strongly associated genes from TWAS and FCT were ascertained for each trait and tissue. Genes were selected for further analysis if they were identified by more than one test of phenotype-genotype associations, i.e. they overlapped the top hit for several analyses/tissues (e.g. overlapping top hits for both GWAS and TWAS leaf). The Arabidopsis orthologs of these genes were collected with INPARANOID (Remm et al., 2001) and used for GO term enrichment analysis in biological function (GO Ontology database DOI: 10.5281/zenodo.4081749 Released 2020-10-09) (Ashburner et al., 2000; Carbon et al., 2019; Mi et al., 2019). PANTHER overrepresentation test was used with Fisher’s test and FDR-adjusted *p*-values, with significance declared at α<0.05. GO biological processes significantly and >2.5-fold enriched were further considered, and the genes contained within these GO categories were considered the strongest candidates to underlie variation in stomatal conductance traits.

## Supplementary material

Supplementary Table S1: LD blocks containing the top 0.1% SNPs from GWAS. The chromosome, upper and lower threshold (bp) for each LD block is also given.

Supplementary Table S2: Top 0.1% strongest GWAS results for each trait, including SNP chromosome and position, marker R^2^ and minor allele effect size.

Supplementary Table S3: TWAS results from each tissue and trait, including chromosome position and R^2^ for each gene.

Supplementary Table S4: FCT results from each tissue and trait, including the GWAS and TWAS *p*-values taken from Supplementary Tables S2-3 and used to calculate an FCT *p*-value for each gene.

Supplementary Table S5: Summary of genes appearing in the top results for GWAS, TWAS leaf, TWAS shoot, FCT leaf and FCT shoot. An “x” identifies genes present in the top associations for an analysis/tissue. The number of analyses that a gene overlapped the top hits for is also shown, ranging from 1 to 5, where n. overlaps=1 indicates a gene was in the top hits for a single analysis/tissue for a given trait, and n. overlaps=5 indicates a gene was in the top hits for all analyses/tissues for a given trait. The Arabidopsis ortholog of each gene is also given. Arabidopsis orthologs of genes that had n. overlaps of at least 2 were further investigated by GO enrichment analysis (Supplementary Table S6). Genes that were present in a GO enrichment category that was significantly and >2.5-fold enriched are identified.

Supplementary Table S6: GO enrichment analysis of Arabidopsis orthologs of genes overlapping top hits for multiple analyses/tissues. Significantly enriched GO biological processes are shown by hierarchical clustering, with the broadest categories further to the right and categories nested within them listed further to the left. Arrows show hierarchical relationships between GO terms. The n. of known genes in Arabidopsis corresponding to each GO term is given, as well as the observed and expected n. of genes, the fold-enrichment (i.e. ratio of observed to expected genes), and *p*-value used to determine whether the enrichment was statistically significant.

Supplementary Table S7: Summary of the most promising candidate genes, selected because they belong to a GO biological process category significantly enriched by >2.5-fold among the subset of genes overlapping the top results for GWAS, TWAS leaf, TWAS shoot, FCT leaf, and/or FCT shoot. Because GO terms are nested, the broadest GO biological process and its fold-enrichment is presented, along with its nested sub-categories that were also significantly and >2.5-fold enriched. Descriptions for each Arabidopsis gene are given. Genes are identified that were determined by a literature survey to be implicated in a WUE-related trait: the reference used to make this determination is also given. Full references are in supplementary material S1.

Supplementary Table S8: BLUPs of all traits for all accessions. Supplementary Table S9: Full GWAS results for each trait.

Supplementary Material S1: Full reference list from literature review of genes in Supplementary Table S7.

Supplementary Figure S1: Pearson’s correlation coefficients (*r*, top right panels), pairwise correlation scatterplots (bottom left panels) and density plots (diagonal panels) for BLUPs of stomatal traits. Data are the same as in Figure 5.

Supplementary Figure S2: GWAS, TWAS and FCT results for *g_s light_*. A: Upset plot showing the number of overlapping genes between the top hits in GWAS, TWAS leaf, TWAS shoot, FCT leaf, and/or FCT shoot. B: Manhattan plots of GWAS, TWAS and FCT results.

Supplementary Figure S3: GWAS, TWAS and FCT results for *g_s shade_*. A: Upset plot showing the number of overlapping genes between the top hits in GWAS, TWAS leaf, TWAS shoot, FCT leaf, and/or FCT shoot. B: Manhattan plots of GWAS, TWAS and FCT results.

Supplementary Figure S4: GWAS, TWAS and FCT results for *g_s oscillation max_*. A: Upset plot showing the number of overlapping genes between the top hits in GWAS, TWAS leaf, TWAS shoot, FCT leaf, and/or FCT shoot. B: Manhattan plots of GWAS, TWAS and FCT results.

Supplementary Figure S5: GWAS, TWAS and FCT results for *g_s initial min_*. A: Upset plot showing the number of overlapping genes between the top hits in GWAS, TWAS leaf, TWAS shoot, FCT leaf, and/or FCT shoot. B: Manhattan plots of GWAS, TWAS and FCT results.

Supplementary Figure S6: GWAS, TWAS and FCT results for *g_s Σ shade_*. A: Upset plot showing the number of overlapping genes between the top hits in GWAS, TWAS leaf, TWAS shoot, FCT leaf, and/or FCT shoot. B: Manhattan plots of GWAS, TWAS and FCT results.

Supplementary Figure S7: GWAS, TWAS and FCT results for *g_s light_* - *g_s shade_*. A: Upset plot showing the number of overlapping genes between the top hits in GWAS, TWAS leaf, TWAS shoot, FCT leaf, and/or FCT shoot. B: Manhattan plots of GWAS, TWAS and FCT results.

Supplementary Figure S8: GWAS, TWAS and FCT results for *t_initial min_*. A: Upset plot showing the number of overlapping genes between the top hits in GWAS, TWAS leaf, TWAS shoot, FCT leaf, and/or FCT shoot. B: Manhattan plots of GWAS, TWAS and FCT results.

Supplementary Figure S9: GWAS, TWAS and FCT results for *V_oscillation_*. A: Upset plot showing the number of overlapping genes between the top hits in GWAS, TWAS leaf, TWAS shoot, FCT leaf, and/or FCT shoot. B: Manhattan plots of GWAS, TWAS and FCT results.

Supplementary Figure S10: GWAS, TWAS and FCT results for *g_s oscillation max_* - *g_s oscillation min_*. A: Upset plot showing the number of overlapping genes between the top hits in GWAS, TWAS leaf, TWAS shoot, FCT leaf, and/or FCT shoot. B: Manhattan plots of GWAS, TWAS and FCT results.

Supplementary Figure S11: GWAS, TWAS and FCT results for G2. A: Upset plot showing the number of overlapping genes between the top hits in GWAS, TWAS leaf, TWAS shoot, FCT leaf, and/or FCT shoot. B: Manhattan plots of GWAS, TWAS and FCT results.

Supplementary Figure S12: GWAS, TWAS and FCT results for G1. A: Upset plot showing the number of overlapping genes between the top hits in GWAS, TWAS leaf, TWAS shoot, FCT leaf, and/or FCT shoot. B: Manhattan plots of GWAS, TWAS and FCT results.

Supplementary Figure S13: Photograph (right) and corresponding thermal image (left) of experimental setup.

